# Whole-genome resequencing and pan-transcriptome reconstruction highlight the impact of genomic structural variation on secondary metabolism gene clusters in the grapevine Esca pathogen *Phaeoacremonium minimum*

**DOI:** 10.1101/252221

**Authors:** Mélanie Massonnet, Abraham Morales-Cruz, Andrea Minio, Rosa Figueroa-Balderas, Daniel P. Lawrence, Renaud Travadon, Philippe E. Rolshausen, Kendra Baumgartner, Dario Cantu

**Author notes:** Authors contributed equally to this work (Mélanie Massonnet, Abraham Morales-Cruz). Correspondence: Dario Cantu.

## Abstract

The Ascomycete fungus *Phaeoacremonium minimum* is one of the primary causal agents of Esca, a widespread and damaging grapevine trunk disease. Variation in virulence among *Pm. minimum* isolates has been reported, but the underlying genetic basis of the phenotypic variability remains unknown. The goal of this study was to characterize intraspecific genetic diversity and explore its potential impact on virulence functions associated with secondary metabolism, cellular transport, and cell wall decomposition. We generated a chromosome-scale genome assembly, using single molecule real-time sequencing, and resequenced the genomes and transcriptomes of multiple isolates to identify sequence and structural polymorphisms. Numerous insertion and deletion events were found for a total of about 1 Mbp in each isolate. Structural variation in this extremely gene dense genome frequently caused presence/absence polymorphisms of multiple adjacent genes, mostly belonging to biosynthetic clusters associated with secondary metabolism. Because of the observed intraspecific diversity in gene content due to structural variation we concluded that a transcriptome reference developed from a single isolate is insufficient to represent the virulence factor repertoire of the species. We therefore compiled a pan-transcriptome reference of *Pm. minimum* comprising a non-redundant set of 15,245 protein-coding sequences. Using naturally infected field samples expressing Esca symptoms, we demonstrated that mapping of meta-transcriptomics data on a multi-species reference that included the *Pm. minimum* pan-transcriptome allows the profiling of an expanded set of virulence factors, including variable genes associated with secondary metabolism and cellular transport.

## INTRODUCTION

Grapevine trunk diseases (Esca, and Botryosphaeria-, Eutypa-, and Phomopsis-diebacks) are a significant threat to viticulture worldwide (Gramaje *et al*., 2018). They are caused by fungal pathogens that colonize the woody organs of grapevines and, by progressively damaging the vascular tissue, reduce yield and shorten the life span of the infected plant (Kaplan *et al*., 2016). Esca is one of the most destructive trunk diseases (Larignon *et al*., 2009; Mostert *et al*., 2006). Its symptoms include an interveinal yellowing and scorching of leaves (‘tiger-stripe’, **Figure 1A**), delayed bud break and dieback of shoot tips, formation of black spots on berries (‘measles’, **Figure 1B**), black lines or spots in the wood (**Figure 1C**), and, in severe cases, sudden wilting and collapse of the whole plant, also known as vine ‘apoplexy’ (Gubler *et al*. 2015; Mugnai *et al*. 1999; Surico *et al*. 2008).

**Figure 1:**
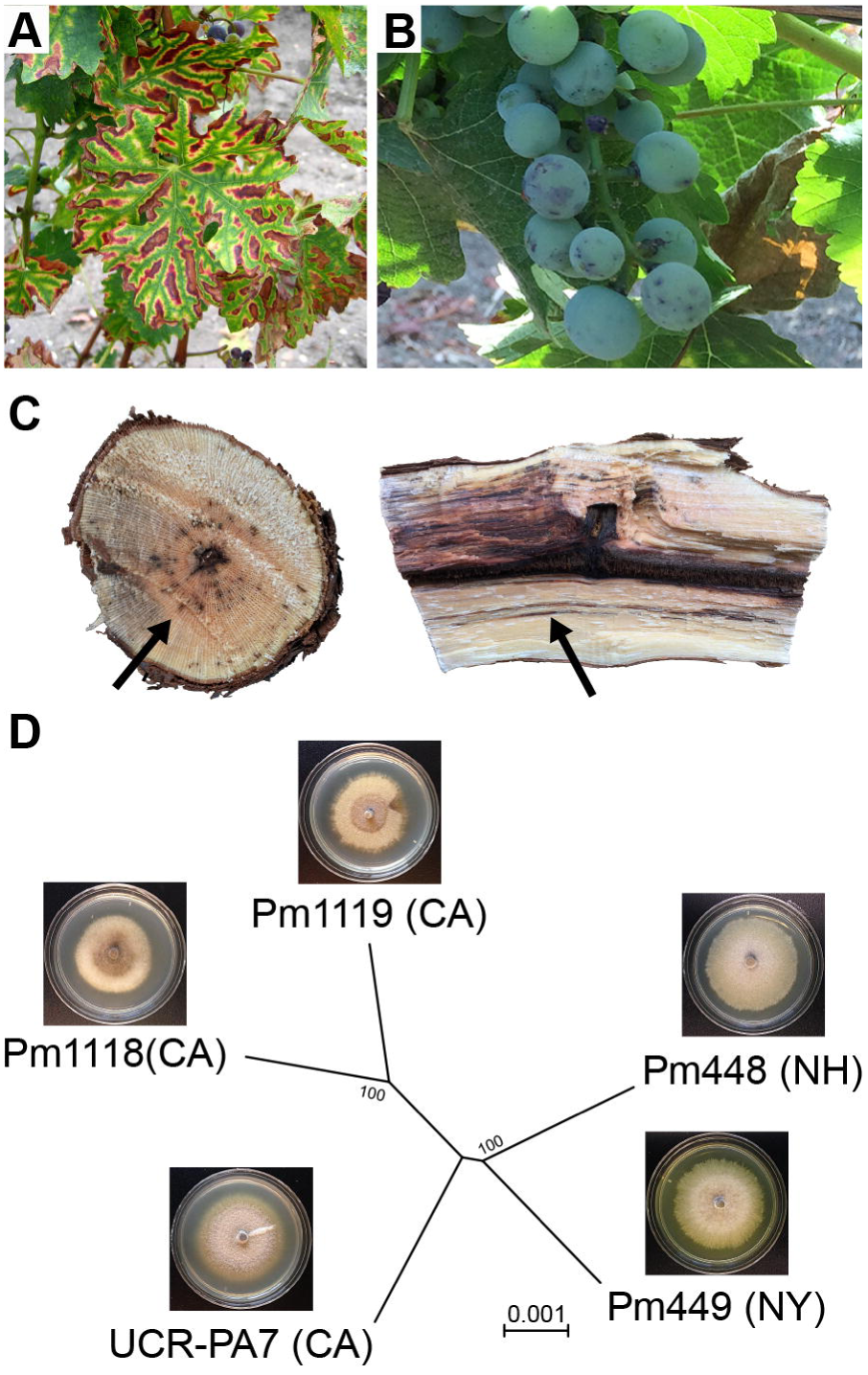
Esca symptoms in grapevine and phylogenetic relation of the five *Pm. minimum* isolates used in this study. (**A**) Typical foliar symptoms of Esca in a red grape cultivar, (**B**) berry spotting (*measles*) and (**C**) black streaking (*arrows*) caused by wood colonization of Esca pathogens. (**D**) Dendrogram illustrating that Pm1119 and Pm448 clustered with Pm1118 and Pm449, respectively, thus reflecting their geographic origins. The dendrogram was constructed with MEGA7 (Kumar *et al*., 2016) using the Neighbor-Joining method (Saitou and Nei, 1987) and was based on a total of 22,242,282 positions. Bootstrap confidence values (100 replicates) are shown next to the branches (Felsenstein, 1985). The evolutionary distances were computed using the Maximum Composite Likelihood method (Tamura *et al*., 2004) and are in the units of the number of base substitutions per site. The tree was visualized with FigTree v.1.4.3 (http://tree.bio.ed.ac.uk/software/figtree). Insets show images of *in vitro* cultures of the five isolates.

Esca is caused by a complex of fungal species, among which are the Ascomycetes *Phaeoacremonium minimum* and *Phaeomoniella chlamydospora* and Basidiomycetes, such as *Fomitiporia mediterranea* (Cloete *et al*., 2014; Fischer, 2006; Surico *et al*., 2008). Esca symptoms are thought to be due to the combined activities of phytotoxic metabolites and cell wall-degrading proteins secreted by the pathogens (Andolfi *et al*., 2011; Mugnai *et al*., 1999). *Pm. minimum* is known to produce several phytotoxic secondary metabolites, including α-glucans and naphthalenone pentaketides, such as scytalone and isosclerone (Bruno and Sparapano, 2006a,b). In addition to phytotoxins, *Pm. minimum* secretes extracellular enzymes that degrade cell wall polysaccharides, such as xylanase, exo- and endo-ß-1,4-glucanase and ß-glucosidase (Valtaud *et al*., 2009). Previous analyses of a draft genome assembly of *Pm. minimum* provided a glimpse of the large number and broad diversity of genes involved in secondary metabolism and cell wall degradation (Blanco-Ulate *et al*., 2013; Morales-Cruz *et al*., 2015). Gene families of these putative virulence factors have undergone distinctive patterns of expansion and contraction in *Pm. minimum* and another Esca pathogen *Ph. chlamydospora*, relative to the genomes of other trunk pathogens, which may explain the differences between Esca symptoms and those of the dieback-type trunk diseases (Morales-Cruz *et al*., 2015).

Significant variability in virulence is reported among *Pm. minimum* isolates (Billones-Baaijens *et al*., 2013; Gramaje *et al*., 2013; Pathrose *et al*., 2014; Pitt *et al*., 2013). This phenotypic variability may reflect the considerable genetic variation at the population level in *Pm. minimum*, which has been described both at the vineyard scale (Borie *et al*., 2002; Péros *et al*., 2000; Tegli *et al*., 2000) and between distant grape regions (Cottral *et al*., 2001; Martín and Martín, 2010; Gramaje *et al*., 2013). Genetic variation in *Pm. minimum* is likely due to its heterothallic reproductive system (Rooney-Latham *et al*., 2005). Indeed, sexual fruiting structures (perithecia) are produced in nature and sexual spores (ascospores) may be important for long-distance dispersal. *Pm. minimum* can also reproduce asexually via production of asexual spores (conidia), which may increase mutation rates, and thus genetic variation, as seen in conidiating lineages of the heterothallic Ascomycete fungus *Neurospora* (Nygren *et al*., 2011).

The impact of this genetic variation on *Pm. minimum* virulence functions remains unknown. In fungal pathogens, single nucleotide polymorphisms (SNPs) and chromosomal structural rearrangements have been shown to underlie gains in pathogenicity, virulence, or adaptation to new environments (Möller and Stukenbrock, 2017). SNPs, for example, may contribute to the generation and maintenance of allelic diversity, which characterizes patterns of host-pathogen co-evolution (Genissel *et al*., 2017; Karasov *et al*., 2014). Structural variations, such as insertions, deletions, and inversions, contribute to phenotypic variation and adaptation by modification of gene dosage, gene expression, or disruption of genes that span boundaries of structural rearrangements (Chow *et al*., 2012; Chuma *et al*., 2011; Jones *et al*., 2014; Qutob *et al*., 2009). For example, subtelomeric tandem duplications yielded a dramatic copy number increase of an arsenite efflux transporter conferring arsenite tolerance in *Cryptococcus neoformans* (Chow *et al*., 2012). Similarly, *Erysiphe necator* populations evolved increased fungicide tolerance to triazole fungicides as a result of multiple duplications of the *Cyp51* gene (Jones *et al*., 2014). Gene duplication and interchromosomal DNA exchange could also lead to formation of novel gene clusters, which may provide an adaptive advantage, as in the case of the *DAL* cluster in yeast (Wong and Wolfe, 2005).

In this study, we investigated the impact that structural variants have on putative virulence functions in *Pm. minimum*. We assembled a chromosome-scale and complete genome of a *Pm. minimum* isolate and resequenced at high-coverage the whole genomes of four additional isolates. We also sequenced the RNA of all isolates grown under different culture conditions, to generate a comprehensive representation of their transcriptomes and expression dynamics. Comparative genome and transcriptome analyses enabled identification of extensive structural variation. Deletions and insertions, in this remarkably dense genome, resulted in hundreds of protein-coding genes that were not shared among isolates. These presence/absence polymorphisms often involved blocks of multiple adjacent virulence factors. Interestingly, the variable fraction of the *Pm. minimum* genome was enriched in clusters associated with secondary metabolism, suggesting that acquisition or loss of secondary metabolism functions may have an adaptive effect on fitness. Finally, we incorporated all core and variable transcripts into a pan-transcriptome, which provided a more comprehensive representation of the virulence repertoire of the species when used as reference for meta-transcriptomic analysis of naturally occurring *Pm. minimum* infections.

## MATERIAL AND METHODS

### Biological material

*Pm. minimum* strains were purified from *V. vinifera* plants (**Data S1: Table S1**) as described in Morales-Cruz *et al*. (2015). For RNAseq, isolates were grown for 28 days in Czapek broth (pH 5.7; (Difco, Detroit, MI, USA)) amended with 0.1% yeast extract (Sigma-Aldrich, Saint-Louis, MO, USA) and 0.1% malt extract (Oxoid Ltd, Basingstoke, UK) at 25°C in both stationary and rotating (150 rpm) conditions in triplicates. Stationary cultures were kept in complete darkness, while rotating cultures were in ambient light.

### DNA extraction, library preparation, sequencing, and assembly

DNA extraction, quality control, and library preparation for PacBio and Illumina sequencing were performed as described in Massonnet *et al*. (2018) and Morales-Cruz *et al*. (2015), respectively. SMRTbell libraries were sequenced using 11 cells of a PacBio RSII system (DNA Technologies Core Facility, University of California Davis), which generated 1,110,178 reads with median and maximum lengths of 8.5 and 50kbp, respectively, for a total of 10.1 Gbp (**Data S1: Table S2**). Illumina sequencing was conducted on a HiSeq2500 sequencing platform in 150 paired-end mode (DNA Technologies Core Facility, University of California Davis), yielding 20,231,286 ± 4,530,073 reads per sample (**Data S1: Table S3**). For UCR-PA7, raw reads were retrieved from NCBI SRA (SRR654175).

Contigs were assembled from PacBio reads with HGAP3.0 and error corrected with Quiver (Chin *et al*., 2013) as described in Massonnet *et al*. (2018). To estimate error rate, Illumina paired-end reads were mapped using Bowtie2 v.2.2.327 (Langmead and Salzberg, 2012), PCR and optical duplicates were removed with Picard tools v.1.119 (http://broadinstitute.github.io/picard/) and sequence variant identified with UnifiedGenotyper from GATK v.3.3.0 (--ploidy 1 --min_base_quality_score 20; McKenna *et al*., 2010). Prior to gene prediction, repetitive regions were masked using a combination of *ab initio* and homology-based approaches, as described in Jones *et al*. (2014). BRAKER prediction was carried out on the soft-masked contigs applying GeneMark-ET with the branch point model. As evidence, we used the paired-end RNA-seq reads retrieved from GSE64404 (Morales-Cruz *et al*., 2015), which were mapped onto the genome assemblies using TopHat v.2.1.0 (Trapnell *et al*., 2009). Only complete protein-coding sequences without internal stop codons were retained. Functional annotations were carried out as described in Massonnet *et al*. (2018).

Illumina reads were trimmed using Trimmomatic v.0.36 (Bolger *et al*., 2014) with options LEADING:3 TRAILING:3 SLIDINGWINDOW:10:20 MINLEN:20 and assembled with SPAdes v.3.10.1 (Bankevich *et al*., 2012) with option --careful. For each genotype, *k-mer* lengths delivering the most contiguous and complete assembly where chosen for the final assembly (**Data S1: Table S3**). Scaffolds (<1 kb) and sequences detected as contaminants by seqclean (Haas *et al*., 2008) were removed. Repeats were masked as described above. Sequences can be retrieved from NCBI (PRJNA421316). Genome sequence of Pm1119, gene prediction and annotation, and pan-transcriptome sequence can be found in **Data S2**. A genome browser of Pm1119 with all relevant tracks can be accessed at https://cantulab.github.io/data.

### Structural variation analysis

Whole genome alignments were performed using NUCmer (MUMmer v3.23; Kurtz *et al*., 2004). SV features and statistics were obtained using dnadiff (Kurtz *et al*., 2004) and assemblytics (Nattestad and Schatz, 2016). SV coordinates were extracted using show-diff. For LUMPY v.0.2.13 (Layer *et al*., 2014) and DELLY2 v.0.7.7 (Rausch *et al.*, 2012), trimmed pair-ended reads were mapped onto Pm1119 using Speedseq v.0.1.2 (Chiang *et al.,* 2015). Only SVs predicted as homozygous alternatives (1/1) in DELLY2 and with at least 4 supporting reads in LUMPY were retained. SVs that overlapped with sites predicted as variant when Pm1119 reads were mapped onto the Pm1119 reference were removed. SV calls of the three methods were compared using bedtools intersect v2.19.1 (Quinlan and Hall, 2010) with a minimum reciprocal overlap of 90% (English *et al.,* 2015). Complete and partial deletions were confirmed by aligning the candidate SV sequences on the respective genome assemblies using GMAP version 2015-11-20 (Wu and Watanabe, 2005).

### Single nucleotide polymorphisms (SNP) calling and phylogeny analysis

Single nucleotide polymorphisms were identified as described above. SNPs were called using the UnifiedGenotyper (GATK v.3.3.0) with the Pm1119 Pacbio assembly as reference. The overall ratio of transition (Tr) over transversion (Tv) mutations was 2.1 ± 0.02. These values are consistent with other studies in fungi (Cantu *et al.,* 2013; Jones *et al.,* 2014) and as expected, are higher than the 0.5 ratio that would be obtained if all substitutions were equally probable. Tr/Tv values were significantly higher in exons (2.7; *P*-value = 9e^-12^) compared to introns (2.0) and intergenic space (1.9; **Data S1: Figure S1**), further supporting the accuracy of gene models and variant calls (DePristo *et al.,* 2011). To identify genes under positive selection we applied the procedure described in Cantu *et al.* (2013). Synthetic sequences incorporating the GATK-detected SNPs were generated using FastaAlternateReferenceMaker of GATK. Orthologous transcripts were then aligned and analyzed using Yn00 (Yang, 2007). Any pair-wise comparisons that yielded a *ω* > 1 were classified as under positive selection.

### RNA extraction, library preparation and sequencing

After 28 days of culture in either stationary or rotating condition, fungal suspensions were vacuum-filtered through 1.6 μm glass microfiber filters (Whatman, Maidstone, UK) and mycelia were collected in a 2-mL micro-centrifuge tube, then immediately frozen in liquid nitrogen and ground to a powder with a TissueLyser II (Qiagen, Hilden, Germany) at 30 Hz for 30 s. One milliliter of TRIzol reagent (Ambion, Austin, TX, USA) was added to the ground mycelia and extraction of total RNA was performed following the manufacturer’s protocol. RNA-seq libraries were prepared using the Illumina TruSeq RNA kit v.2 (Illumina, San Diego, CA, USA) and sequenced on an Illumina HiSeq3000 sequencer (DNA Technologies Core Facility, University of California Davis) in single-end 50-bp mode. Sequences were deposited to Short Read Archive (NCBI; SRA accession: SRP126240; BioProject: PRJNA421316).

### RNAseq, *de novo* transcriptome assembly, identification of isolate-specific transcripts and construction of a pan-transcriptome reference

Reads were first trimmed using Trimmomatic v.0.36 (Bolger *et al.,* 2014) as described above. For each genotype, *de novo* transcriptome assembly was performed using reads from six RNA-seq libraries (3 replicates at stationary + 3 replicates at rotating condition) as input for TRINITY v.2.4.0 (Grabherr *et al.,* 2011). Reconstructed transcripts were then mapped on all genome assemblies using GMAP (Wu and Watanabe, 2005) to determine culture cross-contaminations (**Data S1: Table S4**). We detected significant contamination of Pm448 cultures by Pm449. Consequently, the RNAseq data of Pm448 were not included in further analyses. Transcripts were then mapped with GMAP onto the Pm1119 reference genome to identify variable transcripts (**Data S1: Table S4**). Transcripts that did not map or that mapped with both coverage and identity **≤** 80% were considered not present in the reference. Transcripts derived from mitochondrial genes, with internal stop codon(s), without a stop codon or starting methionine were removed. Transcript redundancies were resolved using the tr2aacds program of Evidential Gene (Gilbert, 2013), which selects from clusters of highly similar contigs the “best” representative transcript based on CDS and protein length. The set of non-redundant transcripts absent in Pm1119 was added to the reference transcriptome to compose the *Pm. minimum* pan-transcriptome. In addition, for each isolate, a private transcriptome was created by removing from the Pm1119 reference transcriptome the transcripts detected as deleted in the isolate and adding the *de novo* assembled complete transcripts detected as not present in Pm1119. Private transcriptomes were then mapped on their own genome assembly using GMAP to determine the genomic coordinates of each transcript (**Data S3**). Co-linearity of the protein-coding genes flanking the locus of insertion was used to identify the orthologous coordinates in the Pm1119 reference genome.

Trimmed single-end reads were mapped onto their corresponding private transcriptome using Bowtie2 v.2.2.6 with parameters: -q -end-to-end -sensitive -no-unal. Then, sam2counts.py v.0.91 (https://github.com/vsbuffalo/sam2counts) was used to extract counts of uniquely mapped reads (Q>30). Details on trimming and mapping results are reported in **Data S1: Table S5**. The Bioconductor package DESeq2 (Love *et al*., 2014) was used for read-count normalization and for statistical testing of differential expression (**Data S4**).

### Closed-reference metatranscriptomics

For meta-transcriptomics, the RNAseq data, retrieved from NCBI SRP092409, consisted of eight libraries from Esca-symptomatic plants, one library from a grapevine with Eutypa dieback-symptoms and one library from a grapevine with no trunk disease symptoms. Reads were quality-trimmed as described above and mapped on a multi-species reference that included the *V. vinifera* cv. ‘PN40024’ transcriptome (v.V1 from http://genomes.cribi.unipd.it/grape/) and the predicted transcriptomes of the 10 fungal species most commonly associated with grapevine trunk diseases as described in Morales-Cruz *et al*. (2017). Concerning *Pm. minimum*, the multi-species reference included the transcriptome of either UCR-PA7 (Morales-Cruz *et al*., 2015), Pm1119, or the *Pm. minimum* pan-transcriptome. Rate of non-specific mapping was evaluated by mapping the six *in vitro* samples of Pm1119 culture onto the meta-reference transcriptome with the *Pm. minimum* pan-transcriptome. Reads mapping onto Pm1119 were randomly subsampled using samtools v.1.3.1 (Li *et al*., 2009) at the median number of reads mapped on *Pm. minimum* pan-transcriptome by the eight Esca samples. Counts of uniquely mapped reads with a mapping quality Q>30 were extracted as described above and details on trimming and mapping results are reported in **Data S5**.

## RESULTS AND DISCUSSION

### Assembly of single molecule real-time sequencing reads generates a complete and highly contiguous reference genome for *Pm. minimum*

The first objective of this study was to generate a complete and highly contiguous genome assembly, to serve as reference for the comparative genome analyses described below. The genome of *Pm. minimum* isolate 1119 (Pm1119, henceforth; **Data S1: Table S1**) was sequenced using single molecule real-time (SMRT) technology at 213x coverage (**Data S1: Table S2**). Sequencing reads were assembled into 25 contigs using HGAP3.0 and error-corrected with Quiver (Chin *et al*., 2013; **Table 1**): 24 contigs formed the nuclear genome with a total size of 47.3 Mbp, whereas the entire mitochondrial genome was assembled into a single 52.5-kbp contig (**Table 1**). N50 and N90 of the nuclear genome were 5.5 and 4.3 Mbp, respectively, representing a significant improvement in sequence contiguity compared to our previous assembly of isolate UCR-PA7, which was generated using short-read sequencing technology (Blanco-Ulate *et al*., 2013; **Data S1: Figure S2**). To evaluate sequence accuracy, we sequenced at 71x coverage the genome of Pm1119 using an Illumina HiSeq2500 system (**Table 1**). Sequence variant analysis with GATK (McKenna *et al*., 2010) detected only 20 single nucleotide sites with discordant base calls between the two technologies. If we assume that Illumina short reads are correct, we could conclude that the contigs generated using SMRT sequencing had a sequence accuracy of 99.999957%.

**Table 1:**
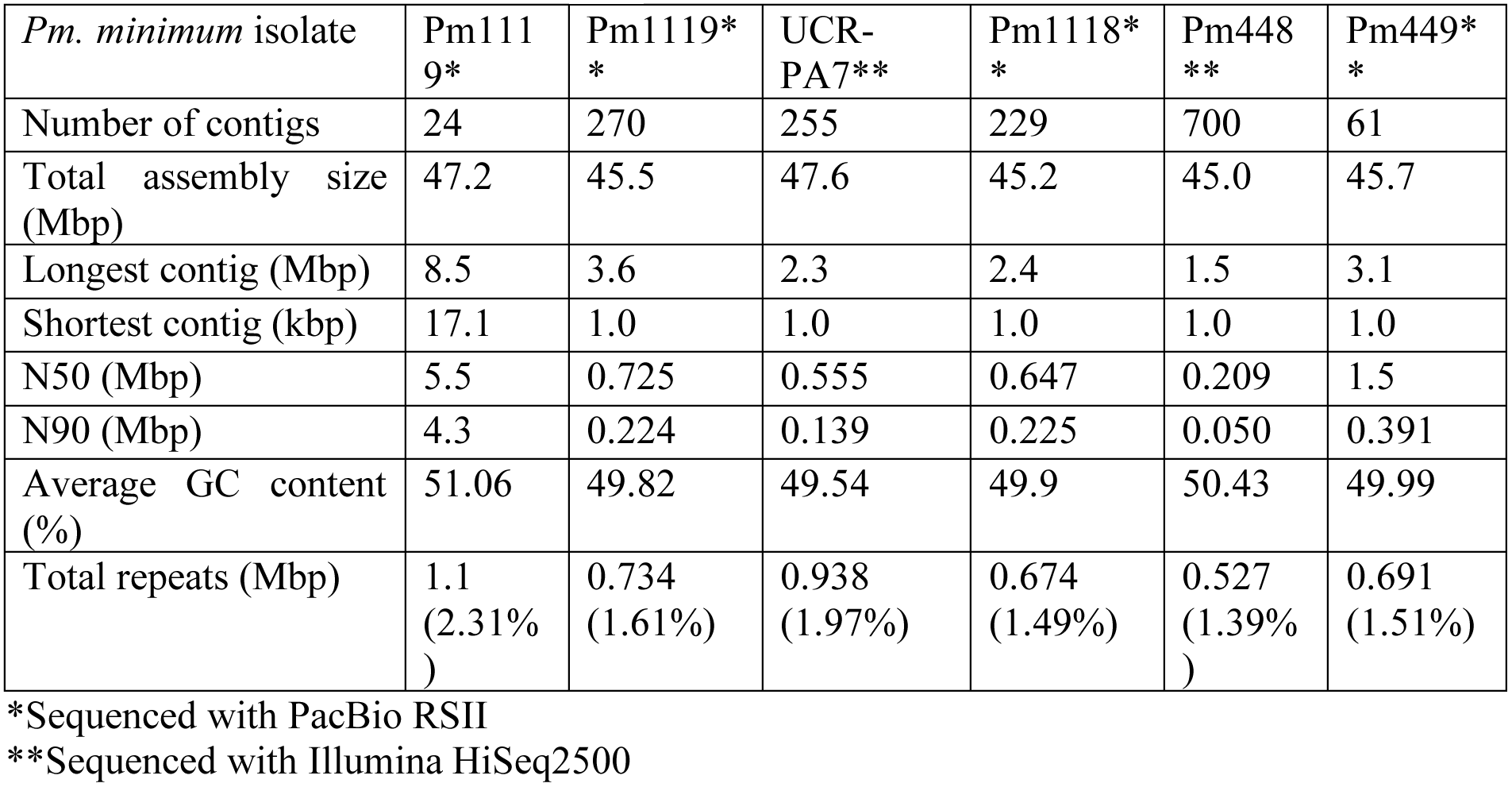
Statistics of the assembled genomes.

The number of chromosomes comprising the *Pm. minimum* nuclear genome is still unknown. In order to determine the degree of fragmentation of the assembly, we searched for the presence of telomeric repeats in the terminal contig sequences. Telomeric repeats (‘5-TTAGGG-3’; Podlevsky *et al*., 2008) were found at both ends of four contigs and at one end of six other contigs, suggesting that at least four chromosomes were assembled telomere-to-telomere (**Data S1: Figure S3**). Protein-coding genes were detected only on nine of the 24 contigs, suggesting that the 15 remaining contigs are fragments derived from intergenic and repetitive regions of the genome, or are potential assembly artifacts. These nine contigs with protein-coding genes comprised 99.2% of the total assembly, with a total size of 46.9 Mbp, which is slightly larger than the genome size estimated using *k*-mer frequency (45.6 Mbp). Approximately 97% of the Core Eukaryotic Genes (Parra *et al*., 2009) and 99.9% of the BUSCO orthologous genes (Simão *et al*., 2015) were found in the assembly, supporting the completeness of the assembled gene space (**Data S1: Table S6**). Only 1.1 Mbp (2.31%) of the Pm1119 genome was composed of interspersed repeats and low complexity DNA sequences (**Table 1**), a repeat content comparable with other grapevine trunk pathogens (3.6 ± 2.0 %; *P*-value = 0.22) but significantly lower than in other Ascomycete plant pathogens (19.8 ± 24.6 %; *P*-value = 0.012; **Data S1: Table S7**). Finally, we compared the assembly with contigs of the same isolate sequenced using short-reads and assembled with SPAdes (**Data S1: Table S3**; Bankevich *et al*., 2012). Only 16 indels, each smaller than 500 bp, for a total of 1,528 bp (**Data S1: Table S8**), were detected with NUCmer (Kurtz *et al*., 2004) validating the overall structural accuracy of the assembly.

Using BRAKER (Hoff *et al*., 2015) and the RNAseq data described below as transcriptional evidence, we identified 14,790 protein-coding genes, including 98.05% of the conserved BUSCO orthologs. Gene density was mostly uniform with 3.4 ± 1.0 genes/10 kbp (**Figure 2**; **Data S1: Figure S4**), a density comparable to other ascomycete plant pathogens (Bindschedler *et al*., 2016). Compared to UCR-PA7, the transcriptome of Pm1119 provided a more comprehensive and accurate representation of the gene space of *Pm. minimum* as shown in **Figure S5** (**Data S1: Table S9**).

**Figure 2:**
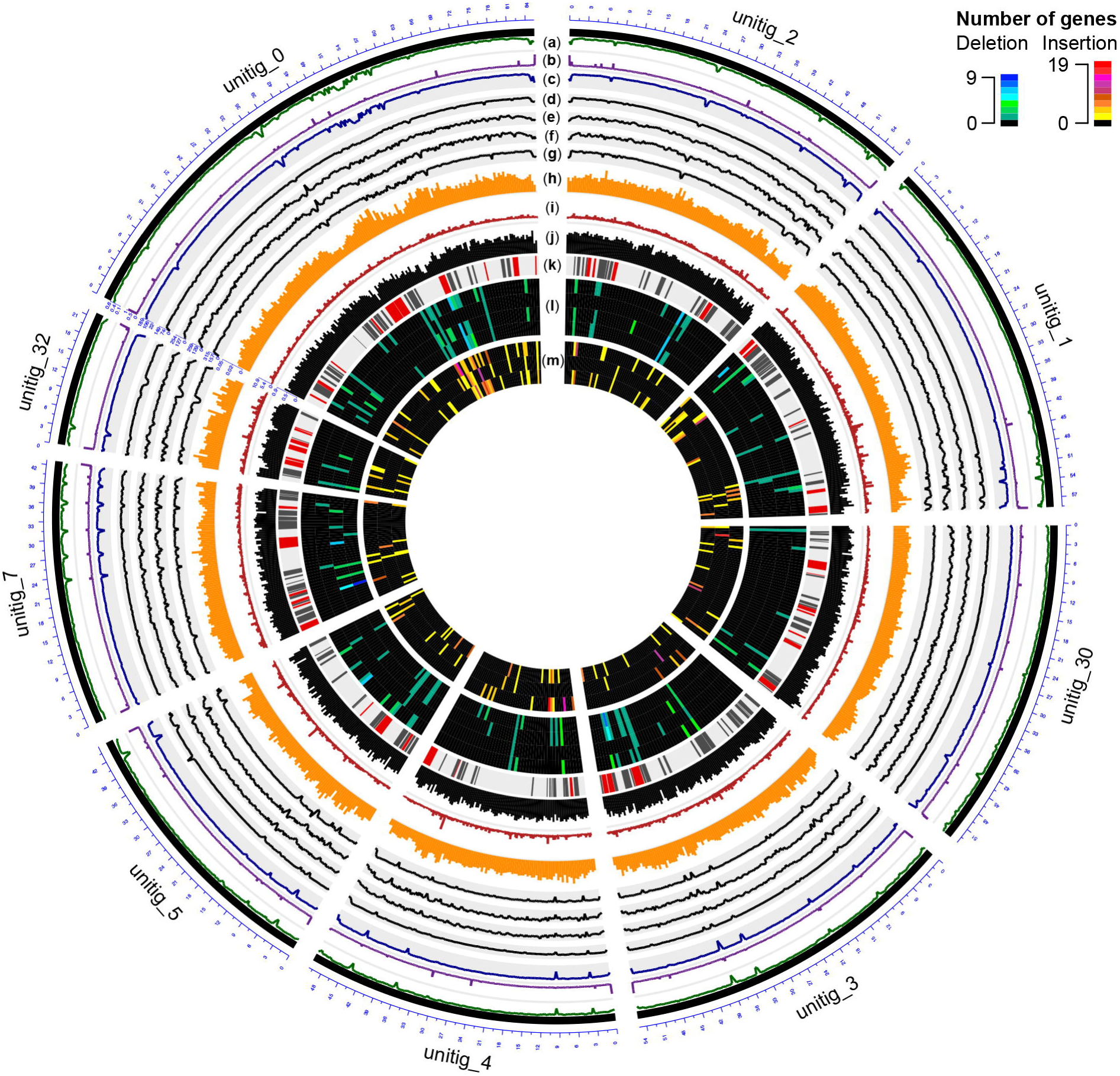
Circular representation of the genome of Pm1119. Different tracks denote: (a) percentage of GC content; (b) repeat density; (c-g) mapping coverage of short Illumina reads: (c) Pm1119, (d) UCR-PA7, (e) Pm1118, (f) Pm448, (g) Pm449; (h) SNP density; (i) omega values; (j) CDS density; (k) BGCs (known classes in red, putative classes in black); (l) Number of genes entirely deleted in UCR-PA7, Pm1118, Pm448, or Pm449; (m) Number of expressed protein-coding genes inserted in UCR-PA7, Pm1118, or Pm449. Figure was prepared using the OmicCircos Bioconductor package (Hu *et al*., 2014).

### Virulence-factor focused annotation shows abundant transport and secondary metabolic functions in the *Pm. minimum* genome

Annotation focused on processes potentially associated with virulence, such as woody-tissue degradation and colonization, cellular transport and secondary metabolism, as described in Morales-Cruz *et al*. (2015). We identified a total of 9,150 genes encoding putative virulence factors, corresponding to 61.9% of *Pm. minimum* predicted transcriptome (**Table 2**; **Data S6**). This set of genes comprised 908 Carbohydrate-Active enZYmes (CAZYmes) including 487 cell wall-degrading enzymes (CWDEs) potentially involved in substrate colonization (**Data S1: Table S10**). Among the set of putative virulence factors were also 52 peroxidases (including two lignin peroxidases), 157 cytochrome P450s (P450s), 2,742 cellular transporters, and 5,712 genes associated with secondary metabolism.

**Table 2:** Number of genes found for each of the major classes of virulence functions in Pm1119 and in the non-redundant set of variable genes identified in the other isolates (*).

|  | Pm1119 transcripts | New transcripts * | Pan-transcriptome |
| --- | --- | --- | --- |
| Protein-coding sequences | 14,790 | 455 | 15,245 |
| Putative virulence factors | 9,150 | 200 | 9,350 |
| CWDEs | 487 | 3 | 490 |
| Peroxidases | 52 | 0 | 52 |
| P450s | 157 | 3 | 160 |
| Cellular transporters | 2,742 | 44 | 2,786 |
| Known BGCs** | 47 (1,739) | 15 (43) | 48 (1,782) |
| Putative BGCs** | 139 (3,973) | 35 (120) | 139 (4,093) |

\*\*The total number of genes in clusters is in parentheses

The annotation of Pm1119 in this study is consistent with the previously observed expansion of families of cellular transporters in *Pm. minimum* and confirmed the relatively smaller set of CAZYmes, compared to the dieback-type pathogens examined in our previous analyses (Morales-Cruz *et al*., 2015). In Pm1119, the Major Facilitator Superfamily (MFS; TCBD code 2.A.1) was the most abundant transporter superfamily, with 816 members and included 200 members of the Sugar Porter (SP) Family (2.A.1.1) and 246 drug-H^+^ antiporter family members [121 DHA1 (2.A.1.2) and 125 DHA2 (2.A.1.3)], which may be involved in toxin secretion (Coleman and Mylonakis, 2009). As observed for other trunk pathogens (Morales-Cruz *et al*., 2015), the genome of *Pm. minimum* comprised a large number of genes potentially involved in secondary metabolism (5,712 genes). These genes are physically grouped on the *Pm. minimum* chromosomes in 186 biosynthetic gene clusters (BGCs), including 47 belonging to known classes, such as polyketide synthesis (PKS), non-ribosomal peptide synthesis (NRPS), and indole, terpene and phosphonate synthesis. The identification of a BGC (BGC_137) involved in phosphonate synthesis is noteworthy considering that some phosphonates are known to have antimicrobial properties. Fungi are known to produce these types of compounds (Wassef and Hendrix, 1976), but the key biosynthetic gene in the BGC (phosphoenolpyruvate phosphonomutase, PEP mutase), has been characterized only in bacteria (Yu *et al*., 2013). Even though one of the predicted proteins of BGC_137 has a putative PEP-mutase domain (BLASTP e-value 6.30e^-60^), until experimentally demonstrated we can only hypothesize that the production of phosphonates may contribute to *Pm. minimum* fitness (Gardner *et al*., 1992; Guest and Grant, 1991). Nonetheless, in the microbiologically complex environment that *Pm. minimum* inhabits [i.e., in mixed infections with other trunk pathogens and non-pathogenic wood-colonizing fungi (Travadon *et al*., 2016), in addition to bacteria (Bruez *et al*., 2015)], it is reasonable to expect this fungus to produce various antimicrobial compounds.

### Comparisons of multiple isolates provides a first assessment of structural variation in the species and its impact on the gene space

To investigate the genomic variability in *Pm. minimum*, we sequenced the genomes of four additional isolates from Esca-symptomatic vines (**Figure 1D**; **Data S1: Table S1**). Strains isolated from distant geographic locations, with distinct colony morphology and *in vitro* growth rates (**Figure 1D**; **Data S1: Figure S6**), were chosen to maximize the potential genetic diversity in the species. An average of 3.4 ± 1.4 Gbp were generated for each isolate, achieving a sequencing coverage of 72 ± 29x (**Data S1: Table S3**). Sequencing reads were directly used to identify single nucleotide polymorphisms (SNPs). Using GATK, we found a total of 1,389,186 SNPs (**Data S1: Table S11**). SNP density was higher in introns (10.8 ± 2.8 SNPs/kbp) compared to exons (4.9 ± 1.5 SNPs/kbp) and intergenic space (8.8 ± 2.4 SNPs/kbp), supporting the overall accuracy of the gene models (**Data S1: Figure S1**). Phylogenetic analysis based on the SNPs (**Figure 1D**) indicated that Pm1118 and Pm448 are genetically closer to Pm1119 and Pm449, respectively. SNP information was used to estimate the selective pressure acting on each of the protein-coding genes in the *Pm. minimum* genome using Yn00 (**Figure 2**; **Data S6**; Li *et al*., 1985; Yang, 2007). Interestingly, gene members of the BGCs involved in terpene synthesis were significantly overrepresented (*P*-value = 2.8e^-3^; **Data S1: Table S12**) among the 2,136 protein-coding genes under positive selection (*ω* > 1), suggesting this pathway may have played an important role in recent adaptation of *Pm. minimum* (Vitti *et al*., 2013).

To explore genomic structural diversity, we assembled the genomes of the four isolates and compared all assemblies (**Table 1**; **Data S1: Table S3**). Total assembly size varied slightly among isolates, from 45 Mbp for Pm448 to 47.6 Mbp for UCR-PA7, and N50 values ranged from 0.2 Mbp for Pm448 to 1.5 Mbp for Pm449. NUCmer analysis of whole-genome alignments (**Data S1: Figure S7**) determined that at least 91.9% of the assemblies aligned to Pm1119 (**Data S1: Table S13**), and identified multiple insertion/deletion events [≥ 50 bp/indel; ∼1 Mbp of structural variant sites (SVs) per isolate] in all genotypes relative to Pm1119 (**Data S1: Table S8**). Because whole-genome alignments depend on the contiguity and completeness of the sequences, the results of NUCmer may have been confounded by the fragmentation of the isolates that were assembled from short-reads (Alkan *et al*., 2011). We therefore also applied LUMPY (Layer *et al*., 2014) and DELLY (Rausch *et al*., 2012), both of which use sequencing read alignment information to identify SVs. Pm1119 was used as reference for both analyses and, therefore, the detected SVs are all relative to Pm1119. LUMPY and DELLY identified 7,133 and 8,355 SVs, respectively. Only 1,233 SVs were identified by both programs. These common SVs included 263 translocations, 861 deletions, 44 duplications and 65 inversions (**Figure 3A**; **Data S7**). Forty six percent of the SVs (570 SVs) identified by both programs were also detected by NUCmer (**Figure 3A**). The limited overlap between results of the three programs confirmed previous reports that showed the importance of using multiple callers to reduce the false discovery rate at the cost of reducing sensitivity of SV detection (Jeffares *et al*., 2017; Sedlazeck *et al*., 2017). All but one of the SVs detected by the three programs were deletions (568 ≥ 50 bp SVs; 1.01 Mb total size; **Data S7**). All three methods identified one interchromosomal translocation in Pm448, whereas they did not agree on any insertion event relative to Pm1119, demonstrating the difficulty in detecting this type of structural variation. UCR-PA7, Pm448 and Pm449 presented on average 228 ± 1 deletions corresponding to 479 ± 25 kbp (**Table 3**), while Pm1118 showed fewer events (166) for a shorter total length of 256 kbp, confirming that Pm1118 is genetically closer to Pm1119 (**Figure 1D**).

**Figure 3:**
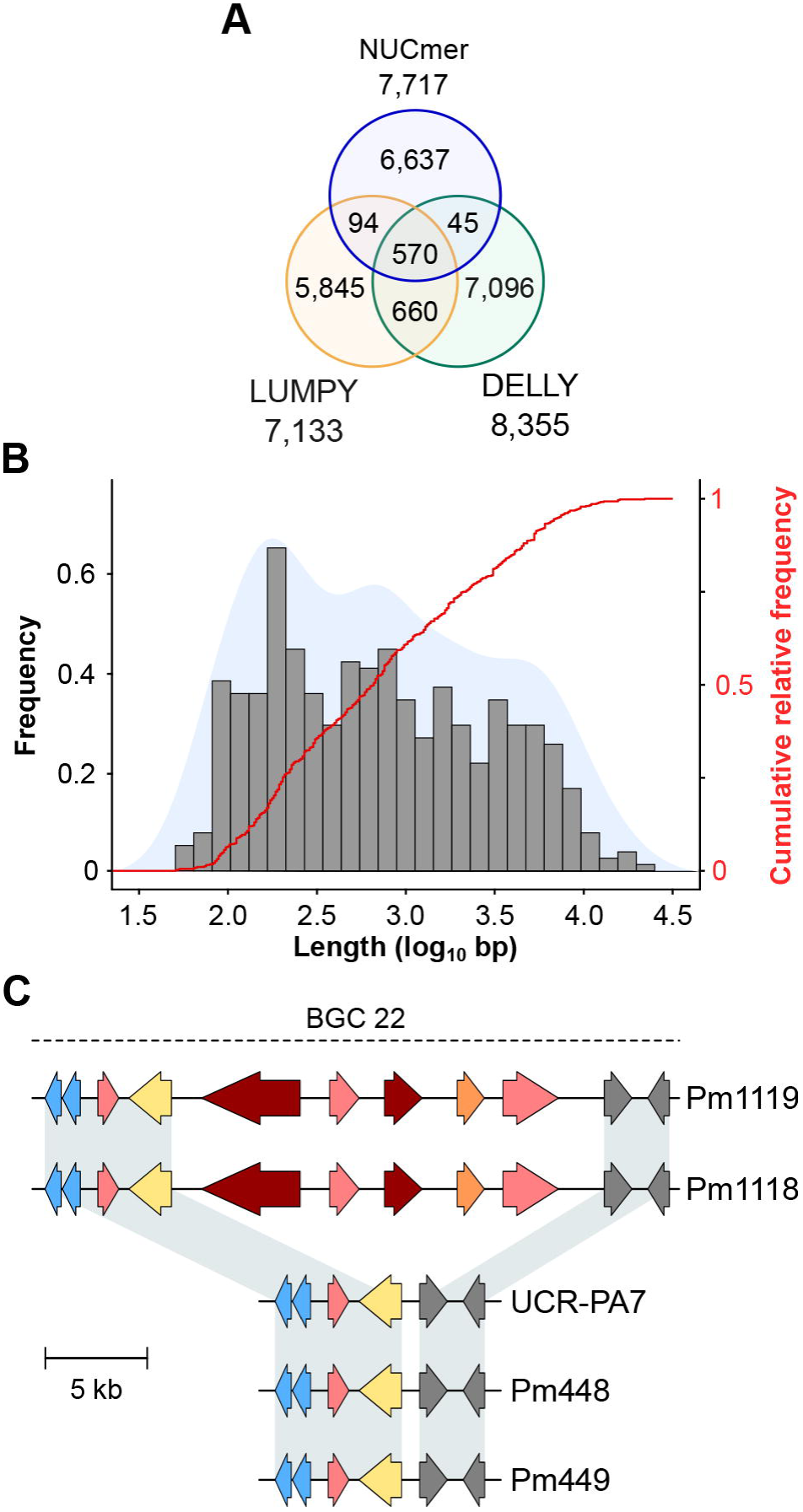
(**A**) Venn diagrams showing the overlap between results of the three methods used to call structural variants (SVs) sites. (**B**) Size distribution of the deleted genomic regions in the four isolates compared to Pm1119. (**C**) Example of a deletion event occurring in UCR-PA7, Pm448 and Pm449 relative to Pm1118 and Pm1119, which affected the composition of a BGC associated with polyketide synthesis. Arrows represent genes coding for core biosynthetic genes (red), additional biosynthetic genes (pink), P450s (blue), cellular transporters (yellow) and FAD-binding proteins (orange). Grey arrows correspond to genes predicted to be part of the biosynthetic gene cluster (BGC), but with other annotations.

**Table 3:** Size, number, and composition of the structural variants identified when comparing the four *Pm. minimum* isolates with Pm1119. Indel genes belonging to BGCs were tested for overrepresentation using Fisher’s exact test. *P*-values are indicated. SV, structural variant. Del, deletions; Ins, Insertions. N.S., statistically non-significant.

|  | UCR-PA7 |  |  | Pm1118 |  |  | Pm448 |  |  | Pm449 |  |  |
| --- | --- | --- | --- | --- | --- | --- | --- | --- | --- | --- | --- | --- |
| SVs | Total | Del | Ins | Total | Del | Ins | Total | Del | Ins | Total | Del | Ins |
| Number of SV | 308 | 227 | 81 | 211 | 166 | 45 | 227 | 227 | 0 | 311 | 229 | 82 |
| Total SV size (kbp) | 630.5 | 471.2 | 159.3 | 341.8 | 256.3 | 85.4 | 508.1 | 508.1 | 0 | 599.1 | 458.5 | 141.4 |
| Number of genes in SVs | 249 | 86 | 163 | 123 | 30 | 93 | 91 | 91 | 0 | 208 | 80 | 128 |
| Genes members of BGCs in SVs | 98 | 51 | 47 | 48 | 11 | 37 | 53 | 53 | 0 | 93 | 50 | 43 |
| BGC members enrichment (*P*-value) | N.S. | 0.000069 | N.S. | N.S. | N.S. | N.S. | 0.000099 | 0.000099 | N/A | N.S. | 0.000011 | N.S. |

Comparison of deletion events among isolates (**Data S1: Figure S8A**) revealed that few events were shared by the four isolates (19/568) and the majority of deletions were isolate-specific (390/568). Pm448 and Pm449 shared almost half of their deletions (105), reflecting their close genetic relationship (**Figure 1D**). The size of deletions ranged from 51 bp to 22 kbp, with a median size of 663 bp (**Figure 3B**). As expected in genomes with a very dense gene space, deletions led to the removal of several protein-coding genes in the four isolates, relative to Pm1119 (**Table 3**; **Figure 4**; **Data S1: Figure S8B**). Interestingly, the detected SVs often encompassed regions in the genome encoding putative virulence functions, such as secondary metabolism and cell wall degradation. Entirely-deleted genes in UCR-PA7, Pm448 and Pm449 were significantly enriched in genes belonging to BGCs (*P*-value ≤ 0.01; **Figure 4**; **Data S8**). Genes involved in polyketide synthesis (t1pks) were significantly overrepresented among entirely- and partially-deleted genes in UCR-PA7 and Pm448, whereas two deletion events resulted in the removal of six of the 30 genes belonging to the BGC involved in phosphonate synthesis in Pm449. We also identified a deletion in UCR-PA7, Pm448 and Pm449 that included five adjacent genes all belonging to BGC_22, which is potentially involved in polyketide synthesis (**Figure 3C**). The genes affected by the indel encode a polyketide synthase and an halogenase (the two core biosynthetic enzymes of the BGC), as well as two *O*-methyltransferases and a FAD-binding monooxygenase, which may be involved in chemical modifications of the final polyketide. Deletions were also enriched (*P*-value ≤ 0.01) in genes involved in cell wall degradation, with the partial removal of two genes encoding enzymes involved in hemicellulose degradation (CE3s; **Data S8**).

**Figure 4:**
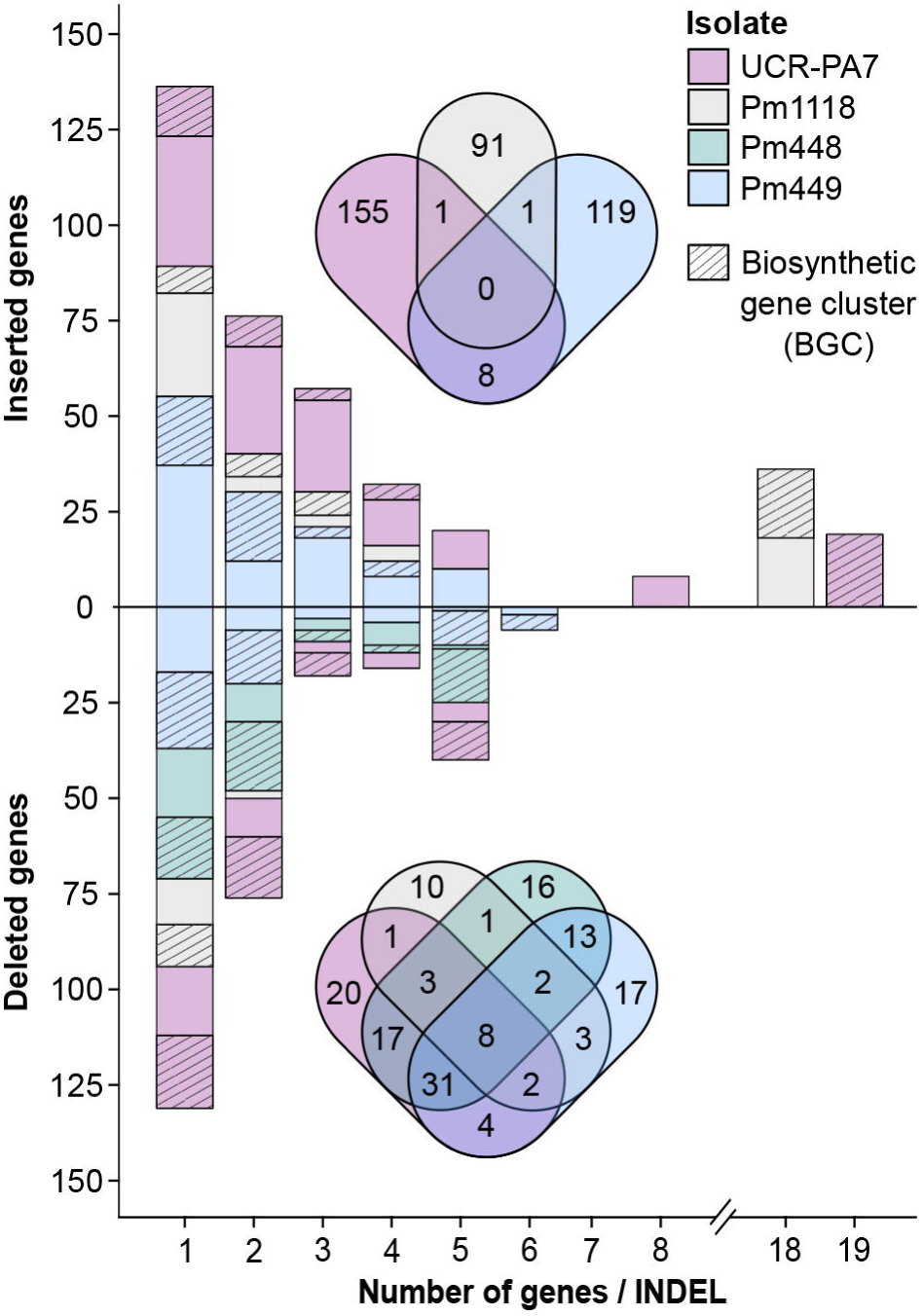
Size and amount of detected indel events encompassing protein-coding genes. The bar plot shows the number of genes and the size of the gene clusters in structural variant sites as well as the proportion of genes associated with biosynthetic gene clusters (BGCs). Venn diagrams show the overlap between genes in structural variant sites detected in the different *Pm. minimum* isolates.

Our findings of structural variation in physically clustered putative virulence factors and, furthermore, significant overrepresentation of BGCs among the variable sites suggest that acquisition or loss of secondary metabolism functions may have an adaptive effect on fitness in *Pm. minimum*. While the acquisition of BGCs may contribute to virulence or antimicrobial activities (Slot, 2017), the loss of accessory products of the secondary metabolism may be adaptive, for example, by evading recognition of the plant immune system (Raffaele and Kamoun, 2012). Patterns of presence/absence polymorphisms of virulence genes have been identified in other populations of fungal pathogens, mainly those with a biotrophic lifestyle (Dai *et al*., 2010; Faino *et al*., 2016; Gout *et al*., 2007; Plissonneau *et al*., 2016; Sharma *et al*., 2014).

### Comparison of *de novo* assembled transcriptomes identifies additional indel events and variable genes in *Pm. minimum*

The analysis of structural variation described above failed to identify any insertion event relative to the reference genome. To identify variable genes that are not presented in the reference, we therefore used an alternative approach: direct comparisons of protein-coding sequences of each isolate with the gene space of the reference genome. This approach has previously identified variable genes in plants (Hansey *et al*., 2012; Hirsch *et al*., 2014; Jin *et al*., 2016). Due to the potential bias caused by the fragmentation of the genomic assemblies of the resequenced isolates, we compared the transcriptomes reconstructed by *de novo* assembly of high-coverage RNA sequencing (RNA-seq) reads. To maximize the diversity and completeness of the sequenced transcriptomes, all isolates were cultured *in vitro,* to generate a higher transcript coverage compared to *in planta* samples (Massonnet *et al*., 2018). Both stationary and rotating cultures were used, to increase the number of protein-coding genes expressed under different culture conditions known to affect both fungal growth and gene expression (Feng and Leonard, 1998; Ibrahim *et al*., 2015; Moreno *et al*., 2007; **Data S1: Table S5**). The transcriptome of each isolate was *de novo* assembled by pooling the reads obtained from three replicates per culture condition. An average of 25,833 ± 5,970 transcripts per isolate were assembled using Trinity (Grabherr *et al*., 2011; **Data S1: Table S4**). The contigs were then mapped on Pm1119 to identify transcripts absent from the reference genome. All of the *de novo* assembled transcripts of Pm1119 mapped onto the Pm1119 genome, thereby confirming the completeness of the gene space in the reference. The transcripts from UCR-PA7, Pm1118 and Pm449 that did not map onto the Pm1119 genome were merged using EvidentialGene (Gilbert, 2013), to generate a non-redundant set of protein-coding sequences (CDS). We identified a total of 455 CDS encoding complete proteins that were not present in the Pm1119 reference: 11 of these were shared by two isolates, whereas 195, 98, and 151 were found only in UCR-PA7, Pm1118 and Pm449, respectively (**Data S3**; **Data S1: Figure S9**). Predicted proteins of the 455 new transcripts were 349 ± 236 amino acid long, which is comparable to the proteins predicted in Pm1119 (**Data S1: Figure S10**). Three of these predicted proteins were annotated as CAZYmes with plant cell wall-degrading functions, three as P450s, 44 as transporters and 150 as members of BGCs (**Data S3**).

By mapping the 455 CDS on their respective genomes, we identified the coordinates of each insertion relative to Pm1119 (**Data S3**; **Table 3**). Many of the insertions involved blocks of multiple genes: 42%, 24% and 33% were insertions of more than one gene in UCR-PA7, Pm449 and Pm1118, respectively (**Figure 4**). The largest inserted block involved 19 adjacent genes in UCR-PA7. In this isolate, we also identified a single SV that involved a complete BGC associated with terpene synthesis, composed of three adjacent genes encoding a P450, a C6 finger transcription factor, and a terpene cyclase. Interestingly, one third of the indels were flanked at both sides by parts of BGCs, further supporting the hypothesis that BGCs are hotspots for fungal genome evolution (Wisecaver *et al*., 2014).

### Analysis of expression of structural variant gene clusters reveals the impact of indel on co-expression of adjacent genes

Physically clustered genes tend to be co-expressed, due to shared regulatory mechanisms (Lawler *et al*., 2013; Massonnet *et al*., 2018). Therefore, to assess the extent of co-expression in the *Pm. minimum* transcriptome and, further, the impact SVs may have on the co-expression of clustered virulence factors, we analyzed the genome-wide patterns of expression dynamics among isolates. RNAseq reads of each isolate were mapped on their respective transcriptomes constructed by combining the shared CDS with Pm1119 and their private *de novo* assembled CDS, as described above. For each isolate, an average of approximately 6 million reads per sample were mapped, detecting an average of 96.5 ± 0.6% of the CDS per isolate (**Data S1: Table S5**). Over 200 additional transcripts were detected under rotating conditions, the majority of which (56.5 ± 3.3%) were associated with secondary metabolism (**Data S1: Figure S11**). An average of 5,824 ± 2,259 transcripts were detected as differentially expressed between stationary and rotating cultures (*P*-value < 0.05; **Data S4**). More than one third of the differentially expressed genes (DEGs) were members of BGCs, confirming the effect of rotation on secondary metabolism on cultured fungi (**Data S1: Table S14**; Ibrahim *et al*., 2015; Shih *et al*., 2007). Approximately 24% of the DEGs of each isolate were composed of genomic clusters containing at least three adjacent co-expressed genes (**Data S1: Table S15**), confirming that transcriptional modulation in *Pm. minimum* involves groups of physically clustered protein-coding regions, as seen in other trunk pathogens (Massonnet *et al*., 2018). The analysis also showed the transcriptional modulation of a total of 295 genes involved in SVs (74 ± 40 per isolate). Interestingly, some co-expressed genomic clusters contained both genes from the Pm1119 reference genome and genes present only in specific isolates (**Figure 5**), suggesting that structural variation within co-expressed genomic clusters does not affect the co-regulation of the other BGC members. In addition, some groups of co-expressed genes were composed only of genes associated with an individual indel. These included the terpene biosynthetic cluster (BGC_187) identified in UCR-PA7 (**Figure 5**).

**Figure 5:**
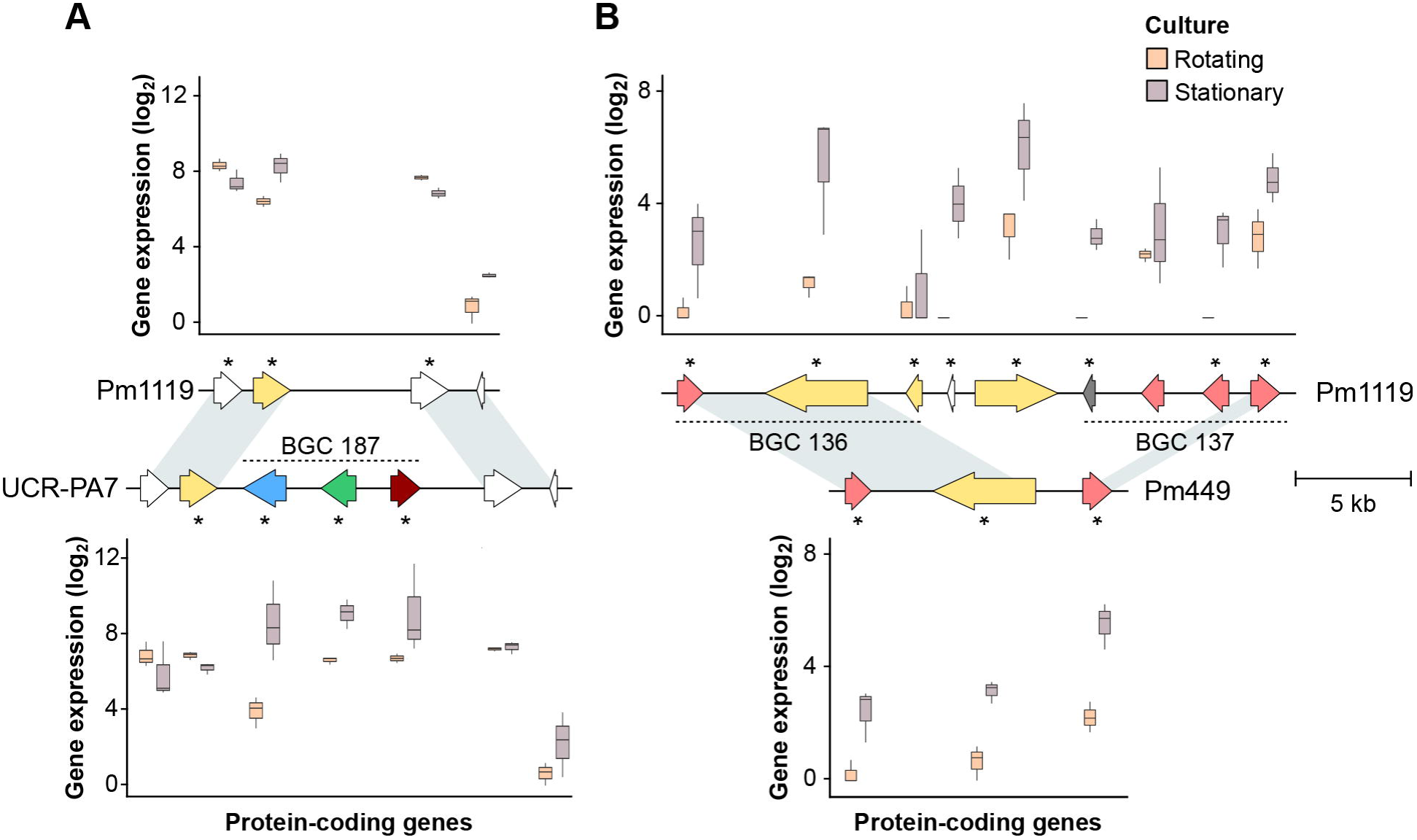
Examples of indels involving co-expressed gene clusters. In (**A**), the whole co-expressed gene cluster BGC_187 identified in UCR-PA7 is deleted in Pm1119. In (**B**), the deletion of adjacent co-expressed genes did not alter the co-expression pattern of the genes flanking the structural variant site. Asterisks identify genes significantly differentially expressed *in vitro* between rotating and stationary cultures. Arrows represent genes coding for core biosynthetic genes (red), additional biosynthetic genes (pink), P450s (blue), cellular transporters (yellow) and transcription factors (green). Grey arrows correspond to genes predicted to be part of the biosynthetic gene cluster (BGC), but with other annotations.

### The addition of the pan-transcriptome to a multi-species reference expands the set of detectable *Pm. minimum* virulence activity in mixed infections in the field

We previously showed that by mapping RNAseq reads on a multi-species reference, comprised of trunk pathogens and common wood-colonizing (non-pathogenic) fungi, we can profile within the mixed infections that naturally occur in the field the expression of putative virulence functions of individual fungi (Morales-Cruz *et al*., 2017). With such a high level of SV involving the gene space and clusters of putative virulence factors, however, we hypothesized that a single genome reference is not sufficient to represent the complete repertoire of virulence functions of *Pm. minimum.* We therefore compiled a transcriptome reference, a pan-transcriptome, which incorporated the variable genes identified in all isolates, i.e. the non-redundant set of CDSs identified in the resequenced isolates. This preliminary pan-transcriptome comprised 14,642 core genes and 603 variable genes. Approximately half of the variable genes were composed of putative virulence factors, mostly associated with secondary metabolism (232 genes) and cellular transport (64 genes). RNA-seq data from the same vine samples we previously examined, collected from Esca-symptomatic vines, were mapped on the following: a multi-species transcriptome that included the genome sequence of grape ‘PN40024’, nine trunk pathogens, and either the CDS of UCR-PA7, the CDS of Pm1119, or the pan-transcriptome of *Pm. minimum*.

The inclusion of the Pm1119 reference resulted in an average increase of 13.4% of the number of reads assigned to *Pm. minimum*, compared to UCR-PA7, without affecting the read counts attributed to the other trunk pathogens. This demonstrates the value of a more complete and contiguous genome in transcriptomic studies (**Figure 6A**; **Data S5**). The inclusion of the pan-transcriptome led to only a slight increase in total read mapping compared to Pm1119, resulting in the detection of 10.6% of the variable CDS (**Figure 6B**). In total, 257 variable transcripts (43% of the variable transcriptome) were detected across the eight vine samples, including 28 transcripts encoding cellular transporters and 94 transcripts associated with secondary metabolism. Significantly fewer genes were mapped when the RNAseq data from the *in vitro* cultures of Pm1119 were aligned onto the same multi-species references, highlighting a low rate of non-specific alignments on variable CDS in the pan-transcriptome. The detection in natural occurring infections of a large portion of the variable transcriptome, and especially of the secondary metabolism-associated variable transcripts, confirms the validity of incorporating pan-transcriptomes in closed-reference metatranscriptomic studies and further suggests that variable genes may play a role during grapevine infections.

**Figure 6:**
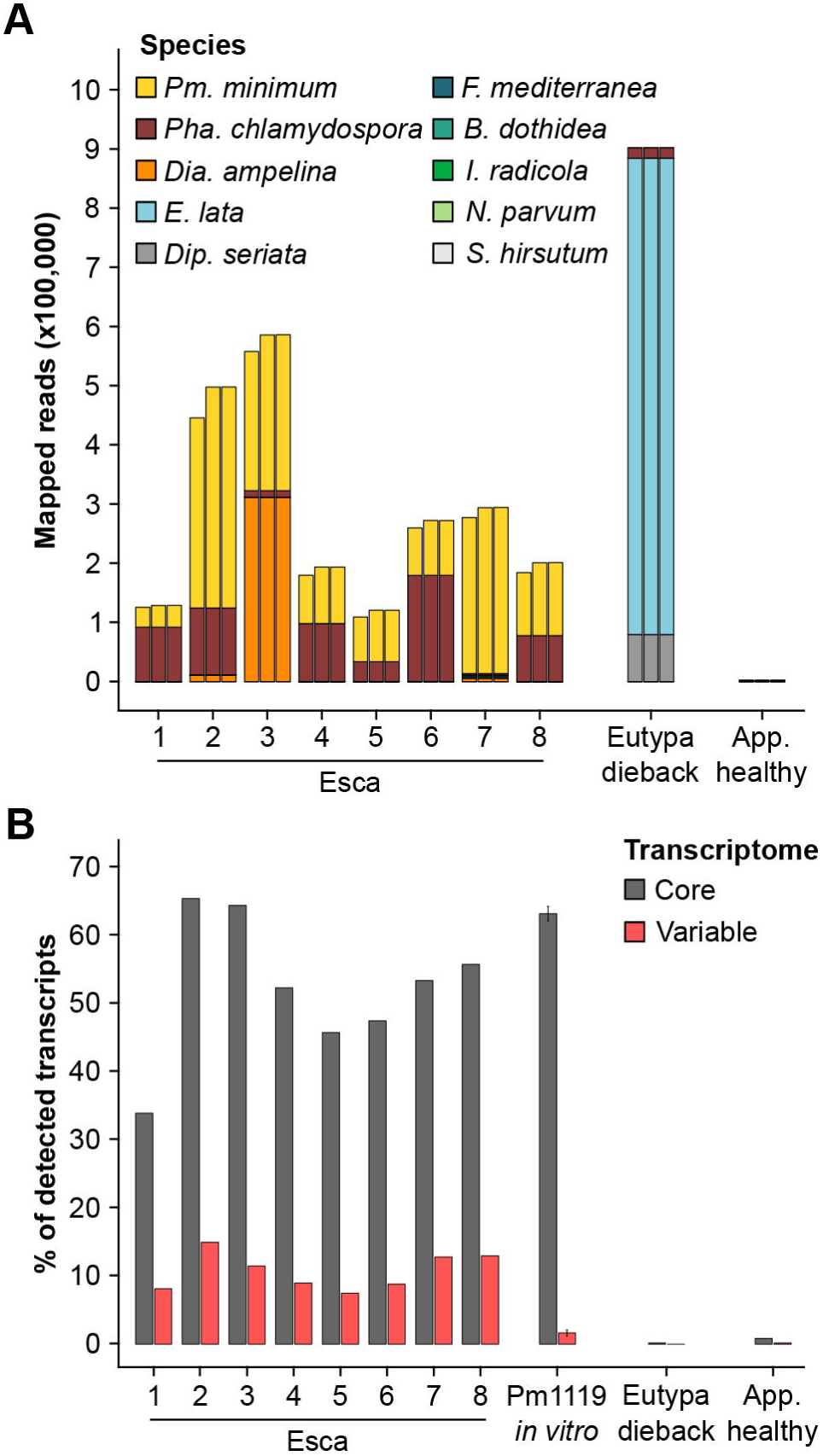
Mapping results of metatranscriptomics data on a multi-species reference including different *Pm. minimum* transcriptome references. (**A**) Stacked bars show the counts of metatransciptomics reads aligned to the transcriptomes of each trunk pathogen included in the multi-species reference using for *Pm. minimum*, from left to right, the transcriptomes of UCR-PA7 (Blanco-Ulate *et al*., 2013), Pm1119, or the pan-transcriptome. (**B**) Percentage of the core and variable transcripts detected in each sample when using *Pm. minimum* pan-transcriptome in the multi-species reference transcriptome. RNAseq data from Pm1119 *in vitro* cultures were included to determine the level of non-specific mapping on variable genes. *Pm*., *Phaeoacremonium*; *Pha*., *Phaeomoniella*; *Dia*., *Diaporthe*; *E*., *Eutypa*; *Dip*., *Diplodia*; *F*., *Fomitiporia*; *B*. *Botryosphaeria*; *I*., *Ilyonectria*. App. healthy, apparently healthy plant without trunk-disease symptoms.

### Conclusions

In this study, we described the genomic diversity among isolates of *Pm. minimum* and showed that detectable structural variation impacted blocks of adjacent virulence genes, preferentially those forming biosynthetic gene clusters involved in secondary metabolism. Because in sexually reproducing fungi like *Pm. minimum*, selection pressure is expected to rapidly eliminate deleterious genes or alleles, it is reasonable to hypothesize that the observed structural variation is maintained because it has adaptive effect on fitness. This hypothesis is also supported by the key role that toxins, a product of secondary metabolism, play during plant colonization (Andolfi *et al*., 2011; Kimura *et al*., 2001) and interactions with other microorganisms (Braga *et al*., 2016). However, we cannot rule out the alternative scenario in which variable genes are rare because they have only a marginal deleterious effect on fitness and, therefore, are not easily lost by microbial populations (Vos and Eyre-Walker, 2017). More experiments are required to understand the role that acquisition or loss of the variable functions play in *Pm. minimum* adaptation. As sequencing costs continue to decline, we can expect that genome-wide association studies, based on whole-genome resequencing of hundreds of isolates, will help link structural variation to pathogen virulence. With the ability to now genetically transform *Pm. minimum* (Pierron *et al*., 2015), the addition or deletion of variable genes, combined with the appropriate experiments to assess *Pm. minimum* fitness, will shed light on the evolutionary role played by structural polymorphisms and the associated variable functions.

## Acknowledgments

This work was funded by the USDA, National Institute of Food and Agriculture, Specialty Crop Research Initiative (grant 2012-51181-19954). DC was also supported by the Louis P. Martini Endowment. We thank Albre Brown for the pictures of Esca-symptomatic plants.

## Authors and Contributors

DC, PER and KB conceived the study. DL and RT carried out the culture experiment. RFB and AMC performed the DNA and RNA extraction, and prepared the sequencing libraries. MM, AM, AMC and DC carried out the computational analysis. MM and DC wrote the manuscript.

All authors read and approved the final manuscript.

## Supplementary Material

### Supplementary Data

**Data S1**: Supplementary tables and figures.

**Data S2**: Genome assembly and protein-coding gene coordinates of Pm1119 and pan-transcriptome sequences.

**Data S3**: Functional annotation of the 455 transcripts not present in Pm1119 and genomic location of the shared Pm1119 CDS and the private CDS on the genomes of each isolate.

**Data S4**: Differentially expressed genes between rotating and stationary culture conditions for each isolate (adj. *P*-value < 0.05).

**Data S5**: Result of the metatranscriptomics analysis of field grapevine samples. (**A**) Statistics of raw, trimmed and mapped RNA-seq data. (**B**) Number of total reads aligned on each grapevine trunk pathogen species when using as reference the multispecies reference and for *Pm. minimum* either UCR-PA7 (Blanco-Ulate *et al*., 2013), Pm1119, or the *Pm. minimum* pan-transcriptome. (**C**) Number of detected *Pm. minimum* transcripts. (**D**) List of the 265 *Pm. minimum* variable transcripts detected across the eight Esca-symptomatic plant samples.

**Data S6**: Annotations of the Pm1119 predicted protein-coding genes.

**Data S7**: Structural variations detected by (**A**) NUCmer, (**B**) DELLY and (**C**) LUMPY, and the overlaps between results of (**D**) NUCmer and DELLY, (**E**) NUCmer and LUMPY, (**F**) DELLY and LUMPY, (**G**) NUCmer, DELLY and LUMPY.

**Data S8**: (**A**) Deletion events identified by the three SV-callers (**A**), genes entirely (**B**) and partially (**C**) deleted and their corresponding enriched functional categories (*P*-value < 0.01; **D** and **E**, respectively).

